# Neural signatures of performance feedback in the paced auditory serial addition task (PASAT): an ERP study

**DOI:** 10.1101/2020.04.06.027359

**Authors:** Anja Sommer, Lukas Ziegler, Christian Plewnia

## Abstract

Due to its importance for successful human behavior, research into cognitive control functioning has gained increasing interest. The paced auditory serial addition task (PASAT) has been used to test and train this fundamental function. It is a challenging task, requiring a high cognitive load in a stressful and frustrating environment. Its underlying neural mechanisms, however, are still unclear. To explore the neural signatures of the PASAT and their link to ongoing cognitive processing, feedback locked event-related potentials were derived from healthy participants during an adaptive 2-back version of the PASAT. Larger neural activation after negative feedback was found for feedback related negativity (FRN), P300 and late positive potential (LPP). In early stages of feedback processing (FRN), a larger difference between positive and negative feedback responses was associated with poorer overall performance, whereas this association was inverted for the later stages (P300 and LPP). This indicates stage-dependent associations of neural activation after negative information and cognitive functioning. Conceivably, increased early responses to negative feedback signify distraction whereas higher activity at later stages reflect cognitive control processes to preserve ongoing performance.

## Introduction

In a world full of competing information and sources of distraction, the ability to maintain coordinated and purposeful behavior is essential to sustain goal directed processes. This requires cognitive control, which comprises different cognitive functions including the ability to pay selective attention, ignore distracting information, turn attention away from stimuli when they prove irrelevant, and the ability to store and manipulate internal representations of information [1]. Especially the inhibition of irrelevant but salient information, like emotional stimuli challenges cognitive control [2]. Cognitive control is a key factor for successful human behavior. Therefore, it is not surprising that dysfunctional cognitive control is increasingly recognized as a key feature of various psychiatric disorders. In fact, research shows that in particular patients suffering from depression are prone to a heightened sensitivity towards negative stimuli, which receive more attention and working memory capacity and therefore impede the maintenance of coordinated and purposeful behavior [3]. This ‘negativity bias’ constitutes an important factor for the development and maintenance of depression as well as a central mechanism of recovery via restoration of cognitive control functioning [1]. Consistently, impairment of goal-directed behavior can be observed in healthy participants when cognitive resources are occupied by emotionally salient distractors [4].

A task used to investigate cognitive control is the ‘paced auditory serial addition task’ (PASAT) [5] in which digits are presented auditorily and participants add the current digit to the digit they heard before. In its adaptive version, inter-stimulus intervals (ISI) decrease (increase) when several consecutive trials are correct (incorrect). The PASAT has been used as a cognitive control task in healthy [6 - 9] as well as clinically depressed [10, 11] and at-risk participants [12], for an overview see [13, 14]. These comprehensive data indicate that this task is particularly suitable to investigate and train cognitive control in both, healthy subjects and psychiatric patients. Regarding its specific mechanism, it has been shown that the PASAT induces frustration and negative affect presumably due to receiving continuously feedback on current performance and working at the individual processing speed limit. Furthermore, the negative affective change induced by the PASAT can in turn be correlated with a lower performance [6] implying that task- or feedback-related irritation must be compensated to maintain goal directed behavior. Therefore, the PASAT challenges cognitive control by means of emotional and cognitive responses to feedback information at a high cognitive load. Regarding the neurophysiological implementation of this competence, prefrontal control on limbic areas plays a key role in overcoming the distraction by negative information and to maintain goal directed behavior [6]. Actually, it has been shown that the dorsolateral prefrontal cortex (dlPFC) is critically involved in the PASAT performance [15]. However, more studies are needed for a better understanding of the mechanisms of action underlying the PASAT. Therefore, the goal of the current study is to investigate the highly time dynamic neural signatures of the PASAT and to capture the conflict of competing negative information and ongoing cognitive functioning in the PASAT. Due to their high temporal resolution event related potentials (ERP) are best suited to find such neural signatures of the PASAT performance.

The feedback related negativity (FRN) is a negative deflection peaking between 200-300 ms after stimulus presentation and is an ERP commonly used to investigate feedback [16, 17]. It is larger for negative than positive feedback and maximal at medial frontal electrode sites [18]. Besides of its informative value concerning current task performance [19, 20] it has been suggested that the FRN indicates the emotional impact of a negative expectation violation [21], implying that feedback does involve emotional processing that captures cognitive resources. Since in our study negative information is operationalized by feedback, we utilized the FRN to investigate early parts of negative information processing. Of note, in our study its registration conditions however differ from the common investigations of the FRN or feedback processing because it summarizes the processing of new information (next digit) and performance feedback (last digit).

Attention allocation to task relevant as well as subsequent memory processes is reflected by the P300. It is a positive deflection peaking between 300-400 ms following stimulus presentation which is maximal at midline-parietal sites [22, 23]. Furthermore, an enhanced amplitude for emotional compared to neutral stimuli can be observed for both, positive and negative content [24 - 26] probably reflecting high inherent motivational salience of emotional stimuli per se. Therefore, it seems to be best suited to study negative information processing during a demanding cognitive task. Moreover, several studies have linked larger P300 amplitudes with performance gains in non-emotional [27, 28] as well as in emotional tasks [29] making it a suitable to investigate associations of neural feedback processing and PASAT performance in our study.

The LPP is known to capture attention allocation toward emotional salient stimuli [30, 31]. It is recorded at centro-parietal sites and begins as early as 200-300 ms post-stimulus. In contrast to the P300 it can outlast the stimulus presentation well beyond several seconds [32]. Therefore, besides its sensitivity to automatic attention allocation to emotional stimuli, it reflects continued processing of emotional content and is regulated by top-down mechanisms. Moreover, the magnitude of the LPP amplitude has also been linked to task performance [33 - 35]. We want to utilize the LPP in our study to capture late neural reactions to negative information in the form of feedback and moreover to investigate its associations with the PASAT performance.

Taken together, with this study, we investigate the time dynamic neural signatures (FRN, P300 and LPP) of the PASAT and aim for a better understanding of its underlying mechanisms. As negative feedback is associated with negative affect and competes with ongoing performance, we assume to find larger amplitudes for negative than positive feedback in all stages of feedback processing. Furthermore, we explore associations of PASAT performance and ERP magnitudes and assume to find significant correlations of cognitive control functioning and neural activation.

## Material and Methods

### Subjects

Twenty-five healthy participants were recruited via internet advertisement. All participants had normal or corrected to normal vision and normal hearing. Exclusion criteria were current psychiatric disorders, neurological disorders, major head injuries or color blindness. They received a financial compensation or course credit for their participation. All participants gave their written informed consent. One participant had to be excluded due to excessive noise in the electroencephalographic (EEG) data (see Electrophysiological Data Processing). The remaining 24 participants (16 female, age: M = 23.71, SD = 4.06) were included in the analysis. To asses neurophysiological characteristics of the participants, we measured complex attention, motor speed, visual-motor conceptual screening and executive functions with the Trail Making Test (TMT) [36]. As a measure for approximate general intelligence we conducted the Multiple Choice Word Fluency Test (MWT-B) [37]. In addition, we measured participant’s working memory with a short version of a digit span test of the Wechsler Adult Intelligence Scale [38]. In our digit span task participants had to memorize 2-8 digits (two trials per level of difficulty, both were used for the calculation of the score) and repeat them in the same order (digit span same order) or in the reverse order (digit span reverse order). See table 1 for demographic and neuropsychological characteristics of the sample and supporting information S1 file for the data underlying the sample characteristics. The study was approved by the local ethics committee and was conducted in compliance with the Declaration of Helsinki.

**Table 1.** Demographic and neurophysiological characteristics of the participants.

| Characteristic | % | M | SD | Range |
| --- | --- | --- | --- | --- |
| Sex (female) | 66.66 |  |  |  |
| Age |  | 23.71 | 4.06 | 20-37 |
| Level of education |  |  |  |  |
| University entrance diploma (yes) | 95.83 |  |  |  |
| TMT-A (s) |  | 23.12 | 7.55 | 14-51 |
| TMT-B (s) |  | 50.28 | 12.51 | 28-78 |
| Digit span same order |  | 8.33 | 1.95 | 5-12 |
| Digit span reverse order |  | 7.38 | 2.20 | 4-12 |
| MWT-B (IQ) |  | 108.58 | 12.63 | 91-145 |

**Notes.** TMT-A = time to complete the Trail Making Test part A; TMT-B = time to complete the Trail Making Test part B; Digit span same order = number of correct remembered digits in the Digit span same order task; Digit span reverse order = number of correct remembered digits in the Digit span reverse order task; MWT-B IQ = Intelligence Quotient assessed with the MWT-B

### Tasks

The tasks *PASAT* and *color presentation* outlined below were computer-based and implemented using PsychoPy2 v1.80.02 [39, 40]. They were presented on a 17-inch monitor.

### Paced auditory serial addition task (PASAT)

We used a 2-back version of an adaptive Paced Auditory Serial Addition Task (2-back PASAT). Participants sat in front of a monitor (distance: approximately 65cm) and heard digits (1-9, duration of presentation: 433-567 ms) via in ear headphones. The task was to add the current digit to the digit they heard before the last one (2-back). Results were indicated by pressing the corresponding key on a keyboard that was equipped with the response letters 2-18. Feedback was given after each trial simultaneously with the presentation of the new digit by presenting green (red) light after correct (incorrect) responses. In order to make the feedback highly salient the whole monitor was filled with the corresponding feedback color (e.g. 17 inch). The duration of the feedback presentation was 433 ms (matched to the presentation duration of the shortest number). Initially the ISI was set to three seconds. The ISI thereby refers to the time in between presented digits as well as feedback, since it was presented simultaneously. The ISI was decreased (increased) after four consecutively correct (incorrect) trials by 100ms to adapt to the capability of the participants. Therefore, the task maintained difficult but manageable throughout the whole session. The task comprised three blocks á 5 minutes with 30 seconds of break in between. The total number of correct trials was used as the main outcome variable. Because the PASAT puts high demands on WM and processing speed it is challenging to stay focused throughout the task and not getting distracted by the feedback information. Accordingly, low cognitive control would result in fewer consecutive correct responses. Therefore, we calculated the proportion of consecutive correct relative to the overall correct responses (performance stability) as a second outcome variable.

### Control Task ‘color presentation’

Since we aimed to test the differential neural responses to feedback valence as indicated by red and green screen color in contrast to the neural activation to red and green color as such, we conducted a control task ‘color presentation’ (CP). Participants were asked to sit in front of a monitor (distance: approximately 65cm) and perceive red and green light peripheral by keeping their gaze on the keyboard just like they would do while performing the 2-back PASAT. The task consisted of two blocks á 2.5 minutes. Red and green light was presented for 433 ms (as in the 2-back PASAT) in random order with a jittered inter stimulus interval (1500-2500 ms).

In sum, there were four conditions for the calculation of the ERPs: green color after a correct trial in the 2-back PASAT (green feedback), red color after an incorrect trial in the 2-back PASAT (red feedback), green color in the CP and red color in the CP. For the 2-back PASAT, only feedback following a response was used (e.g. trials with red feedback for a missing response were excluded from analysis).

### Electroencephalography recording

The electroencephalogram (EEG) was recorded using an elastic cap (EASYCAP GmbH, Hersching, Germany), the actiCHamp amplifier system with 32 active Ag/AgCl electrodes and the corresponding Brain Vision Recorder system (Brain Products GmbH, Gilching, Germany). EEG was registered from 27 scalp sites (FP1, F7, F3, Fz, F4, F8, FC5, FC1, FCz, FC2, FC6, C3, Cz, C4, CP5, CP1, CPz, CP2, CP6, P7, P3, Pz, P4, P8, O1, Oz, O2). Additionally, an electrooculogram (EOG) was recorded. For horizontal eye movements two electrodes were placed approximately one cm left and right of the eyes. One electrode positioned approximately one cm below the left eye and the Fp1 electrode were used to register vertical eye movements. Furthermore, electrodes were placed on the left and right mastoid. The left mastoid served as the online reference and a forehead electrode as the ground. The online sampling rate was 1000 Hz. Impedances were kept below 10kΩ before initiation of the recording.

### Procedure

The experiment took place in a dimly lit, quiet room. After the participants gave their written informed consent, the EEG electrodes were attached to the scalp. Participants were asked to sit quietly during the EEG recording. The experiment started with the CP. After that, participants carried out the 2-back PASAT. To make sure participants understood the instruction of the task, they completed 30 practice trials, which were excluded from analysis. To control for affective effects of the 2-back PASAT, participants completed the 20 item positive and negative affect schedule (PANAS) [41] immediately before and after the 2-back PASAT. That followed a resting phase of 7 minutes during that heart rate measures were obtained, which are not subject of the current paper.

### Electrophysiological Data Processing

We analyzed the EEG data using the EEGLAB toolbox [42] running on MATLAB 9.2 R2017a (The MathWorks, Natick, MA, USA) and the EEGLAB toolbox ERPLAP [43]. The raw EEG was resampled offline to 250 Hz and re-referenced to an average of the left and right mastoids. Band-pass filters with a low and high cutoff of 0.1 and 35 Hz, and a notch-filter at 50 Hz were applied. Ocular artifacts were removed manually using independent component analysis (JADE algorithm) [44]. Subsequently feedback locked epochs were extracted ranging from −100 to 1000 ms relative to feedback (2-back PASAT) and color (CP) onset respectively. Artifact correction was conducted in the epoched EEG. Epochs containing EEG signals exceeding an amplitude of 65 μV within a 100ms moving window or exceeding -65 - 65 μV within the epoch were considered artifacts and were rejected (using the ERPLAB implemented automated artifact detection). Participants with more than 25% of rejected epochs were excluded from further analysis (n = 1). In the 2-back PASAT on average M = 4.12% of the green feedback trials (SD = 5.50%) and M = 4.60% of the red feedback trials (SD = 5.27%) were rejected. In the CP M = 2.57% of the green color trials (SD = 4.67%) and M = 3.25% of the red color trials (SD = 6.76%) were rejected. Overall, 2610 green and 1245 red feedbacks of the 2-back PASAT and 1796 green and 1771 red color trials were included in the ERP analysis. ERPs were constructed by separately averaging trials in the four conditions (green feedback, red feedback, green color, red color). Subsequently we calculated difference waves: positive feedback = green feedback-green color, negative feedback = red feedback – red color. All further ERP analyses refer to these difference waves (see supporting information Figures S1-6 depicting the raw waveforms and scalp maps separately for the 2-back PASAT and CP).

We chose the electrode sites and time windows to measure the FRN, P300 and LPP according to previous literature. The FRN was defined as the mean amplitude within a time window between 200-300 ms following feedback at Fz [18]. The P300 was scored as the average of three centro-parietal sites (Cz, CPz, Pz). Since according to visual inspection there is a large shift in the P300 waveforms due to the FRN, the P300 was defined as the base-to-peak amplitude as follows: we first calculated the peak amplitude (PA) of the most negative peak between 200-300 ms post feedback presentation separately for positive and negative feedback. Then we calculated the PA of the most positive peak between 300-400 ms post feedback presentation separately for positive and negative feedback (PA P300). Afterwards the difference between the peak and the base was calculated separately for positive and negative feedback (base-to-peak P300 = PA P300 - PA of the negative peak between 200-300 ms post feedback presentation [22, 45, 46]. The LPP was scored as the average of five centro-parietal sites (Cz, CP1, CPz, CP2, Pz) and defined as the mean amplitude within a time window between 400-1000 ms following feedback [32, 47].

### Data Analysis

All statistical analyses were performed using SPSS Statistics for Microsoft Windows (version 24.0). See supporting information S1 file for the data underlying the results. To analyze a differential neural activation to positive vs. negative feedback we performed paired t-tests separately for the FRN, P300 and the LPP. To further analyze associations of the valence-specific neural activation (Δ = negative-positive feedback) and changes in the affect ratings with the 2-back PASAT performance (number of correct trials and performance stability) we calculated bivariate correlation analyses using Pearson’s correlation coefficient. Additionally, it could be assumed that a better 2-back PASAT performance would be associated with fewer incorrect trials and therefore with fewer negative feedback. In turn, the mere difference in the presentation frequency of good vs. bad performers could lead to a differential neural reaction to negative feedback and we would not know if an association of the valence-specific neural activation (Δ = negative-positive feedback) and the 2-back PASAT performance could just occur due to this difference and not due to differences in cognitive control functions. Therefore, we additionally calculated the correlation of the number of incorrect trials and the 2-back PASAT performance (number of correct trials). For all analyses, two-tailed tests were used, and a 0.05 level of significance was employed. Post-hoc paired t-tests were conducted where appropriate.

## Results

### Changes in Affect and Behavioral Data

After the 2-back PASAT, overall affect deteriorated significantly as indicated by the PANAS: positive affect ratings decreased [before: *M* = 29.13, *SD* = 5.06; after: *M* = 26.21, *SD* = 5.67; *t*(23) = 2.41, *p* = .025] and negative affect ratings increased [before: *M* = 13.29, *SD* = 2.93; after: *M* = 20.96, *SD* = 9.51; *t*(23) = −4.86, *p* < .001]. There were no significant correlations of the affect ratings with the 2-back PASAT performance (all *p* ≥ .472). Concerning the 2-back PASAT performance, on average, participants gave 113.04 (*SD* = 31.32) correct, and 52.42 (*SD* = 19.68) incorrect responses with 239.54 (*SD* = 31.45) trials overall (including trials without a response).

### Electrophysiological Data

#### Feedback related negativity

Figure 1 displays the grand average waveform (panel A) of the FRN and the mean voltage distribution across the scalp (panel B) for negative and positive feedback separately (note that higher negative values indicate a larger FRN). The mean amplitude FRN for negative feedback was significantly larger (*M* = −0.817, *SD* = 3.223) than for positive feedback [*M* = 0.846, *SD* = 2.843; *t*_(23)_ = 2.671, *p* = .014]. The correlation analysis for the valence-specific neural activation of the FRN (ΔFRN = negative-positive feedback, e.g. a more negative value indicates that the FRN for negative feedback was larger than for positive feedback), revealed a significant association between the ΔFRN and the number of correct trials in the 2-back PASAT (see Figure 1C, note that for all scatterplots the Y-axis is ordered ascendingly according to the values indicating larger ERPs, e.g. for the FRN values are ordered from positive to negative). A smaller ΔFRN (e.g. more positive ΔFRN) was linked to a larger amount of correct trials over all [*r*_(22)_ = 0.425, *p* = .038]. In addition, we found a significant correlation of the ΔFRN and the performance stability. A smaller ΔFRN (e.g. more positive ΔFRN) was linked to a higher performance stability [*r*_(22)_ = 0.433, *p* = .034].

**FIGURE 1.**
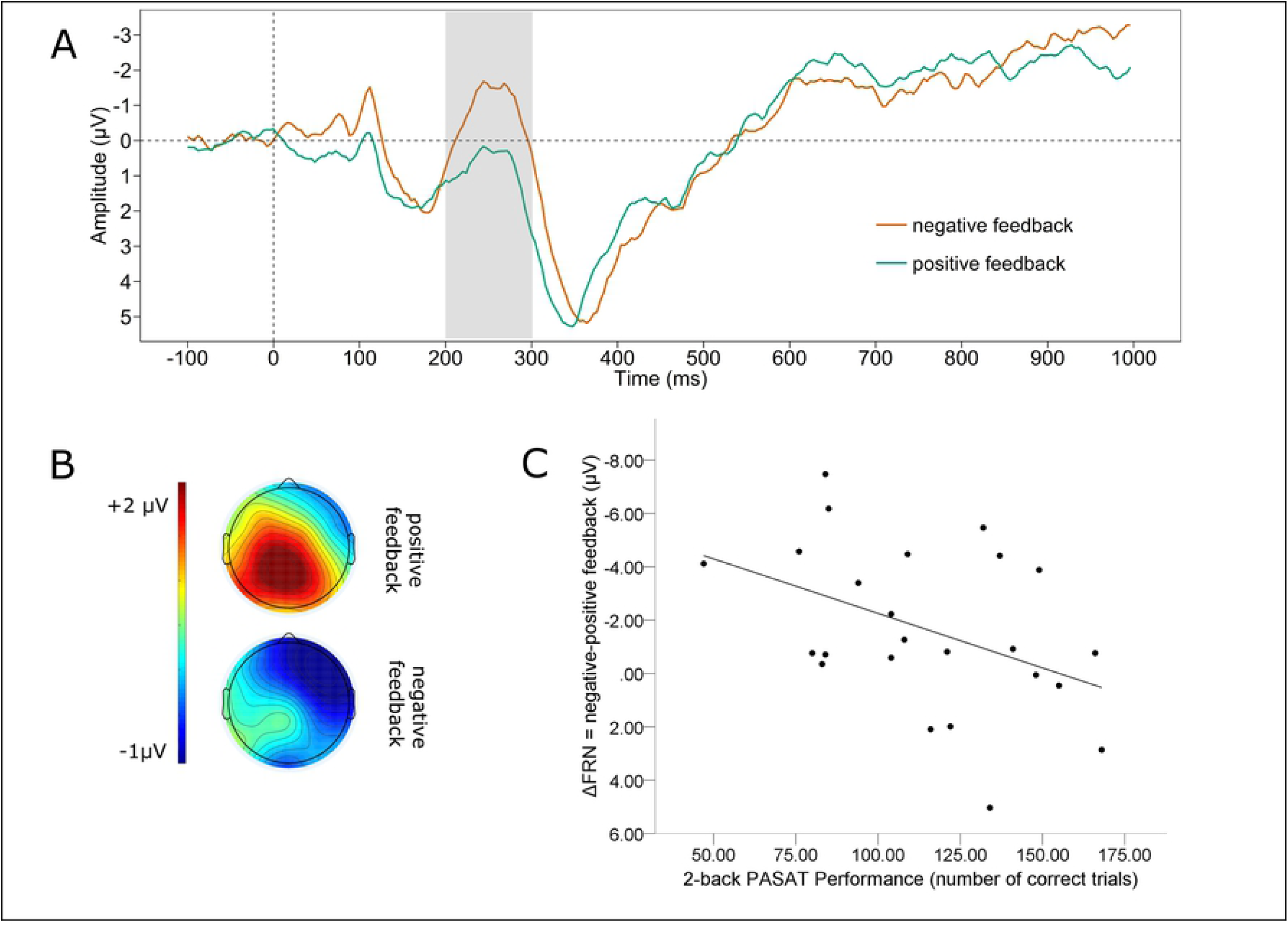
Feedback related negativity. Panel **A** displays the grand average waveform separately for negative and positive feedback at Fz. Panel **B** shows the scalp map displaying the mean voltage distribution for negative and positive feedback separately (200 - 300 ms post feedback). Panel **C** shows a scatterplot displaying the 2-back PASAT performance as a function of the valence-specific FRN (ΔFRN = negative-positive feedback). Note, for the ΔFRN, more negative values indicate a larger amplitude by negative than positive feedback. Therefore, negative values are at the top of the Y-axis.

#### P300

Figure 2 displays the grand average waveform of the P300 (panel A) and the mean voltage distribution across the scalp (panel B) for negative and positive feedback separately. Note that to avoid carry over effects of the shifts in the waveform in the time range of the FRN to the P300, we conducted base-to-peak analyses to define the P300 amplitudes. A paired t-test revealed a significant difference between the P300 for positive and negative feedback. The P300 was significantly larger for negative feedback (*M* = 10.648, *SD* = 4.047) than for positive feedback [*M* = 8.812, *SD* = 3.464; *t*_(23)_ = 3.64, *p* = .001]. For the valence-specific neural activation of the P300 (ΔP300 = negative-positive feedback), we found a significant correlation between the ΔP300 and the number of correct trials in the 2-back PASAT (see Figure 2C). We observed that a larger P300 elicited by negative as compared to positive feedback (e.g. a more positive ΔP300) was linked to more correct trials [*r*_(22)_ = 0.422, *p* = .040]. In addition, we found a significant correlation of the ΔP300 and the performance stability [*r*_(22)_ = 0.465, *p* = .022]. A larger P300 by negative as compared to positive feedback was linked to increases in performance stability.

**FIGURE 2.**
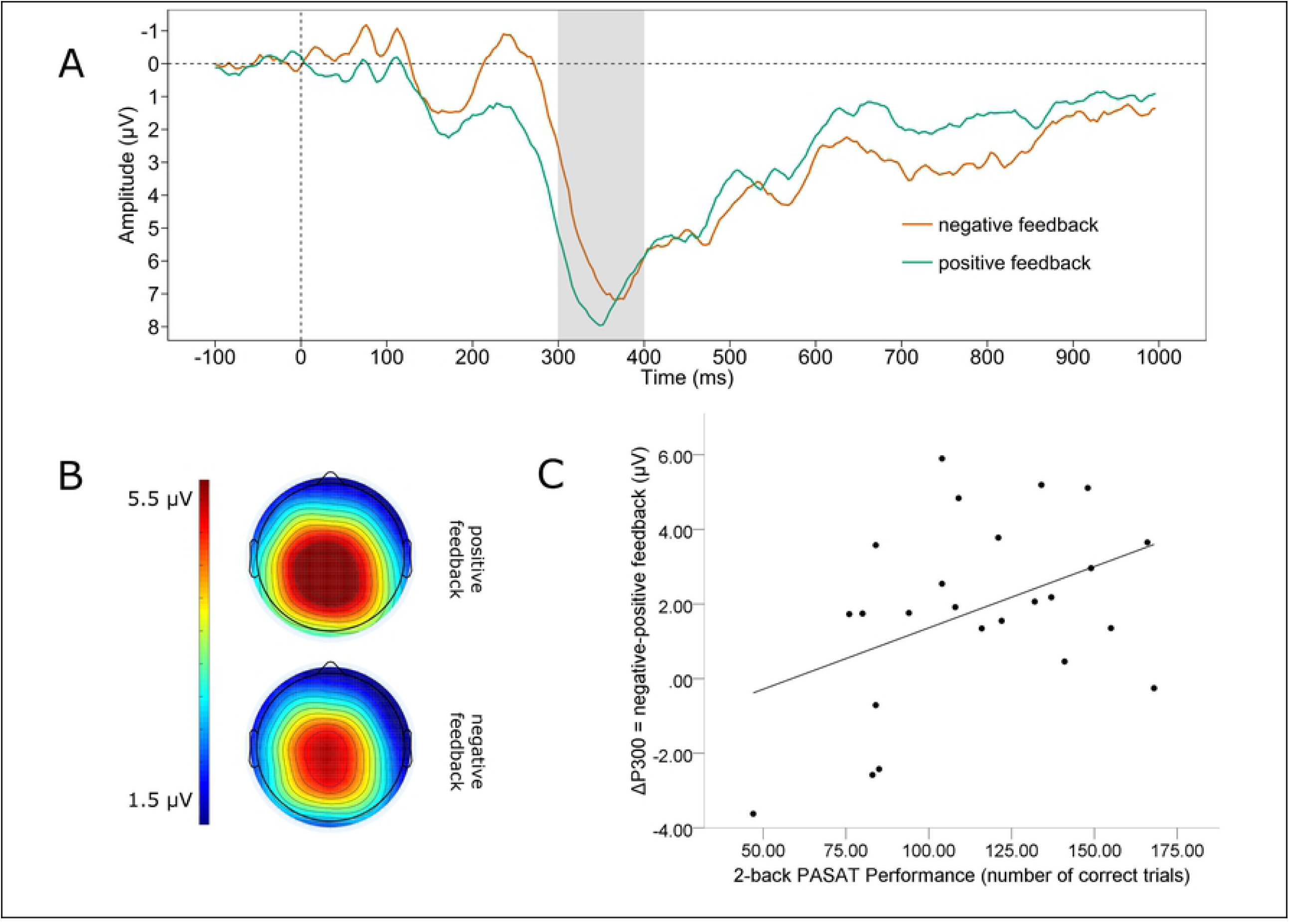
P300. Panel **A** displays the grand average waveform of the P300 separately for negative and positive feedback averaged across Cz, CPz, Pz. Note that a base-to-peak analysis was performed for the P300. Panel **B** shows the scalp map displaying the mean voltage distribution for negative and positive feedback separately (300 - 400 ms post feedback). Panel **C** shows a scatterplot displaying the 2-back PASAT performance as a function of the valence-specific P300 (ΔP300 = negative-positive feedback).

#### Late positive potential

The grand average waveform of the LPP (panel A) and the mean voltage distribution across the scalp (panel B) for negative and positive feedback separately, are depicted in figure 3. We found a significant difference between the mean amplitude LPP for positive and negative feedback. The LPP for negative feedback was significantly larger (M = 3.125, SD = 3.118) compared to positive feedback [M = 2.368, SD = 3.026; t(23) = 2.215, p = .037]. Further, there was a medium effect sized correlation of the valence-specific neural activation of the LPP (ΔLPP = negative-positive feedback) and the number of correct trials, which failed to reach significance [r(22) = 0.300, p = .155]. However, we found a significant correlation of ΔLPP and performance stability in the 2-back PASAT [r(22) = 0.407, p = .049, see Figure 3C]. A larger LPP by negative as compared to positive feedback was linked to increases in performance stability.

**FIGURE 3.**
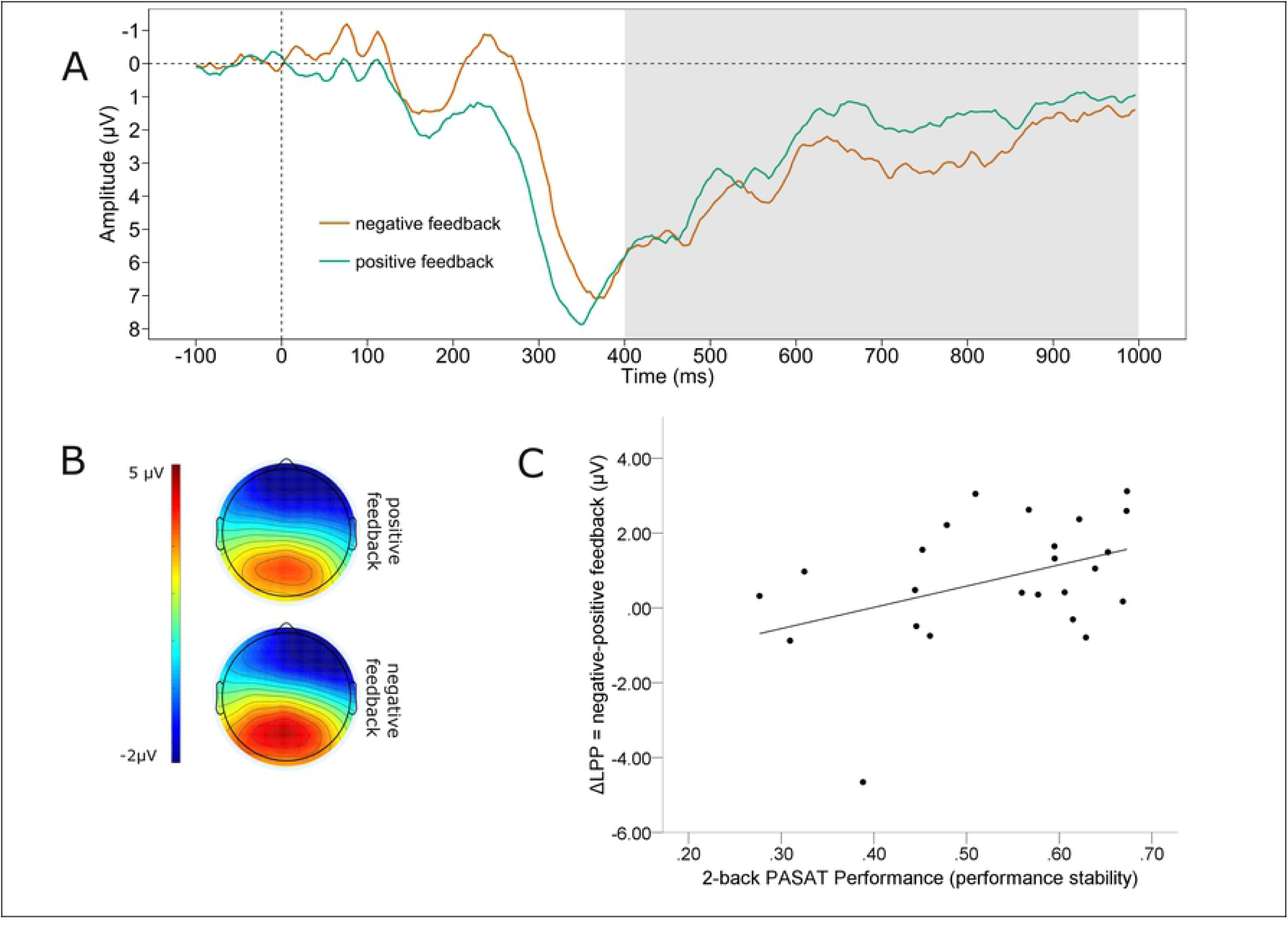
Late positive potential. Panel **A** displays the grand average waveform of the LPP separately for negative and positive feedback averaged across Cz, CPz, Pz, CP1, CP2. Panel **B** shows the scalp map displaying the mean voltage distribution for negative and positive feedback separately (400 - 1000 ms post feedback). Panel **C** shows a scatterplot displaying the 2-back PASAT performance as a function of the valence-specific LPP (ΔLPP = negative-positive feedback).

## Discussion

In this study, we examined electrophysiological characteristics of cognitive control processes and their relation to task performance by means of a challenging and adaptive 2-back PASAT. The main findings are (a) that positive and negative feedback induce a differential neural activation throughout the time course of feedback processing (b) that the valence-specific neural activation (negative-positive feedback) is associated with the 2-back PASAT performance, and (c) that the direction of this association is critically dependent on the stage of feedback processing.

We found that negative information in the form of performance feedback produced similar neural signals for the FRN time range as in common studies investigating feedback processing [48]. Thus, in line with our hypothesis, the FRN was larger for negative feedback than for positive feedback. This observation for the FRN is frequently interpreted as a stronger neural reaction of the anterior cingulate cortex (ACC) for negative than for positive feedback. Since the pMFC including the ACC is known to reflect the motivational value of stimuli [49] this further suggests that in early stages of feedback processing in the 2-back PASAT, negative feedback is probably perceived as more salient than positive feedback. This makes sense, concerning the fact that negative feedback contains important information to adapt behavior according to changing task demands. Therefore, it could be assumed that a more pronounced reaction to errors is beneficial for task performance. In accordance with this assumption several authors describe a beneficial effect of larger FRN and error-related negativity (ERN) amplitudes on task performance [50 - 53]. For example, when learning a sequence of button presses by trial and error the FRN was significantly larger for trials that were followed by a correct response indicating that a larger FRN was associated with a better learning efficacy [54]. However, in our study we found that a larger valence-specific FRN amplitude was associated with poorer task performance, indicating that in the 2-back PASAT the feedback plays a different role compared to common studies investigating the FRN. To understand this, it must be considered that in the present study, besides of its informational value, the negative feedback had also the potential to fundamentally distract from task performance, since it was presented simultaneously with the next target. Therefore, we interpret the FRN as a neural signature of attention allocation towards a distractive negative information as opposed to the task relevant target. This is in line with the assumption that the FRN indicates the emotional impact of negative expectation violation [21]. In accordance with the well-established evidence of a negativity bias linked to a decreased cognitive control in depression, it has been shown that the FRN is enhanced in patients suffering from current as well as remitted depression indicating a hypersensitivity to loss, punishment or negative related stimuli in depression [55 - 57] which reflects reduced cognitive control over emotions. This is consistent with our finding of a poorer 2-back PASAT performance (number of correct trials as well as the performance stability) in healthy participants with larger FRNs. Moreover, this finding suggests that a larger neural activation following negative than positive feedback is linked to an enhanced sensitivity to negative stimuli, which leads to increased distraction, by the valence-specific neural activation in this early stage of feedback processing.

For the P300 we could also confirm our hypothesis of a stronger neural activation for negative than for positive feedback. Consistent with findings showing that the P300 reflects attention allocation towards motivationally and/ or emotionally relevant content, this indicates that negative feedback in the 2-back PASAT is associated with greater resource allocation than positive feedback. This assumption is bolstered by the correlation of the ΔP300 (negative-positive feedback) and the 2-back PASAT: in contrast to the FRN a larger P300 to negative than positive feedback was associated with a larger number of correct trials and a higher performance stability. This finding is in accordance with previous studies showing comparable associations. For instance it has been observed that a better performance in an n-back working memory task was associated with a larger P300 amplitude [27, 28]. Moreover, a larger P300 was found to be associated with more remembered stimuli of emotional content [29]. Therefore, our finding adds further evidence that the additional recruitment of neural activity at this stage of processing leads to performance gains and the maintenance of goal-oriented behavior.

In line with our hypothesis, we also found a larger amplitude for negative than for positive feedback for the LPP. Since a large body of evidence shows that the LPP is larger for emotional than for non-emotional stimuli this indicates that negative feedback was perceived as emotionally more relevant than positive feedback [30]. The fact that negative feedback captures more resources than positive feedback reflected by the LPP suggests that in later stages of feedback processing a negativity bias can be observed. Regarding the valence-specific neural activation of the LPP (ΔLPP = negative-positive feedback) we could observe a similar pattern as for the ΔP300. Although the medium effect sized correlation of the ΔLPP with the number of correct trials in the 2-back PASAT failed to reach significance, we found a significant correlation of the ΔLPP and the performance stability, indicating the same association: a larger LPP by negative than positive feedback was associated with a higher performance stability. Just like the association of the ΔP300 with performance, a stronger neural reaction to negative than positive feedback in later processing stages seems to reflect the recruitment of additional cognitive resources, which increase the effective maintenance of coordinated behavior. Our data are in accordance with results showing a positive relationship between larger LPP amplitudes and task performance. This was observed for example in a delayed working memory task: larger ΔLPPs (negative - positive) evoked by emotional pictures serving as distractors were associated with better task performances [35].

Furthermore, also in an approach avoidance task it was found that larger LPP amplitudes were linked to faster RTs [34]. In contrast, there is also a finding of larger LPP amplitudes associated with performance deteriorations, indicating increased engagement with a distracting stimulus [33]. However, it must be considered that no WM task was used in this study, but a speeded response task, focusing on the investigation of attentional processes and less demanding cognitive functions as opposed to our study. In sum, our results for the LPP and the P300 seem to be in line with the idea of an additional recruitment of cognitive resources by emotional stimuli [58], at least in these late stages of feedback processing.

Consistent with previous studies, we found affect ratings significantly decreased after 2-back PASAT performance [6, 59, 60]. However, there was no significant correlation between the PANAS affect ratings and the 2-back PASAT performance [6]. Nevertheless, their functional association is underlined by the correlation between the evoked potentials indicating emotional processing and task performance. For that matter, the use of self-report questionnaires like the PANAS might be not sufficiently precise to detect latent affect changes.

Taken together it appears that in the early stages of feedback processing (<300 ms following feedback) in the 2-back PASAT, less automatic resource allocation towards negative than positive feedback is beneficial for task performance. Whereas in later stages (>300 ms following feedback) this association is inverted: a more extensive neural recruitment following negative feedback is linked with better performance. Conceivably, in bad performers increased early (< 300 ms) activation after negative feedback interferes with successful memory updating. Apparently, through largely bottom up driven processing, attentional resources are diverted away from target processing and towards distractive negative information, which is reflected by a large neural response to negative feedback.

Accordingly, good performance is associated with the ability to engage top-down control already at early processing stages and maintaining attentional resources to targets and not negative feedback information. Opposingly, in later stages of feedback processing (> 300 ms), large valence-specific amplitudes seem to reflect resource allocation towards goal directed task processing, indicating successful implementation of top-down control. Therefore, good performers are apparently capable of using the feedback information in a top down driven manner to achieve goal directed behavior, reflected by a large valence specific neural activation in late processing stages.

Overall the ERP signatures we found contribute to a better understanding of the neural mechanisms underlying the PASAT and furthermore help to gain better understand why the PASAT is an efficient cognitive control training and could be a promising, innovative treatment option for patients suffering from depression. Our results suggest that poor performance is associated with increased sensitivity to negative information in early processing stages and reduced allocation of cognitive resources in later stages. As stated above, depressed patients depict a hypersensitivity to negative feedback and negative information in general. PASAT training may help to reduce this hypersensitivity by implementing cognitive control strategies in early processing stages to cope with the frustration caused by PASAT. At the same time, these activated cognitive resources could lead to an effective use of the feedback information in later processing stages. This hypothesis is supported by findings showing a critical involvement of the dlPFC in the PASAT performance [15], which in turn is a neuronal structure underlying cognitive control functioning and has been found to be hypoactive in depressed patients [61]. Our results provide useful tools to test such a possible training mechanism and to determine which patients can benefit from a cognitive control training in the long run.

There are some limitations of the current study. It could be assumed that a better performance in the 2-back PASAT would be confounded by fewer incorrect trials and therefore fewer negative feedback. This would indicate that a differential neural reaction to negative vs. positive feedback could be a result of this difference as opposed to be a marker of cognitive control functions. However, due to the adaptive design of the 2-back PASAT, a good performance goes along with a faster stimulus presentation and as a result, participants make more mistakes. This is also reflected by the missing association of the 2-back PASAT performance and the amount of negative feedback: good performers receive as much negative feedback as bad performers. In addition, there have been a lot of misses in the task (e.g. trials without a response) that were excluded from data analysis. Probably also these misses reflect meaningful information. However, during the experiment we observed that the cause for the misses are manifold: participants simply processed the digits not fast enough; sometimes there actually was a response, but it occurred at the same time the feedback was presented (meaning it was not recorded) or sometimes participants zoned out and did not process the stimuli at all for several trials. Unfortunately, we cannot distinguish between these cases afterwards but at the same time it can be assumed that their informative value for cognitive functions and neural responses to feedback are quite different. Therefore, we decided to exclude misses completely from the analysis.

To conclude, by elucidating the neural mechanisms underlying the PASAT performance, we demonstrate that enhanced neural activity in early processing stages of negative feedback indicates a diversion of cognitive resources towards negative information resulting in reduced goal-oriented behavior. In turn, additional allocation of resources after salient negative information as indicated by a higher P300 and LPP is linked with enhanced performance and may thus represent a neural signature of successful cognitive control of distractive negative information. Our results provide the basis for further studies using and investigating the PASAT as an effective cognitive control task. Based on these results, future studies will further elucidate associations and malleability of negative information processing, cognitive performance and mood regulation in sensitive population groups and psychiatric disorders.

## Supporting information

**S1 File. Dataset underlying the findings**. The sav. file contains all data concerning sample characteristic as well as behavioral and electroencephalographic results underlying our findings.

**S1 Figure. Raw FRN for the 2-back PASAT**. The grand average waveform separately for red and green feedback (left panel) and the scalp map displaying the mean voltage distribution for red and green feedback separately (right panel, 200 - 300 ms post feedback).

**S2 Figure. Raw FRN for the control task ‘color presentation’**. The grand average waveform separately for red and green color (left panel) and the scalp map displaying the mean voltage distribution for red and green color separately (right panel, 200 - 300 ms post feedback).

**S3 Figure. Raw P300 for the 2-back PASAT**. The grand average waveform separately for red and green feedback (left panel) and the scalp map displaying the mean voltage distribution for red and green feedback separately (right panel, 300 - 400 ms post feedback).

**S4 Figure. Raw P300 for the control task ‘color presentation’**. The grand average waveform separately for red and green color (left panel) and the scalp map displaying the mean voltage distribution for red and green color separately (right panel, 300 - 400 ms post feedback).

**S5 Figure. Raw LPP for the 2-back PASAT**. The grand average waveform separately for red and green feedback (left panel) and the scalp map displaying the mean voltage distribution for red and green feedback separately (right panel, 400 - 1000 ms post feedback).

**S6 Figure. Raw LPP the control task ‘color presentation’**. The grand average waveform separately for red and green color (left panel) and the scalp map displaying the mean voltage distribution for red and green color separately (right panel, 400 - 1000 ms post feedback).

